# Neural dynamics in superior colliculus of freely moving mice

**DOI:** 10.1101/2025.04.16.648828

**Authors:** Shelby L. Sharp, Jhoseph Shin, Dylan M. Martins, Keaton Jones, Cristopher M. Niell

**Affiliations:** Institute of Neuroscience and Department of Biology, University of Oregon, United States; Graduate Program in Dynamical Neuroscience, University of California, Santa Barbara, United States

## Abstract

Vision is an active process that depends on head and eye movements to explore the visual environment. Superior colliculus (SC) is known for its role in generating these movements, as well as processing visual information, but has not been studied extensively during free movement in complex visual environments. To determine the impact of active vision, we recorded neural activity across the depth of SC during free movement while simultaneously recording eye and head position. We find that superficial SC (sSC) neurons respond to visual input following gaze-shifting saccadic movements, whereas deep SC (dSC) neurons respond to the movements themselves, as demonstrated by their sustained response in darkness. Additionally, we find motor responses in dSC are more correlated to head movements rather than eye movements. Furthermore, we compared sSC gaze shift responses to known gaze shift responses in primary visual cortex (V1), finding similarities in key response types, although the temporal sequences following gaze shifts differ between the regions. Our results demonstrate distinct visual processing differences between SC and V1 as well as highlighting the various roles SC plays during active vision.

**Highlights:**

- We recorded neural activity across the depths of superior colliculus (SC) in freely moving mice while measuring head and eye movements.
- Neurons in mouse SC respond strongly to gaze shifts, and these responses differ between superficial (sSC) and deep (dSC) layers.
- sSC neurons respond primarily to the visual input during a saccadic movement.
- dSC neurons generally represent the head movement more than the eye movement, independent of visual input.
- While sSC gaze shift responses share similarities with V1 there are unique response profiles in SC that suggest differences in the role of visual processing between SC and V1 during free movement.

## Introduction

When analyzing the visual scene, animals often utilize movement of their eyes, head, and body to sample the environment. This combination of movements is essential to active vision, which is necessary for most visually dependent behaviors. The superior colliculus (SC) is a highly conserved midbrain structure known for its involvement in visuo-motor behaviors^1,2^. SC has a laminar structure with superficial layers (sSC) receiving direct retinal input dedicated to visual processing^2–6^ and deeper layers (dSC) responding to multimodal input and producing motor output^7,8^; this organization makes SC primed to analyze aspects of the visual scene and initiate appropriate movements. A large body of research shows that SC is involved in the generation of head and eye movements across species^8–12^.

It is of particular importance to investigate the activity of SC across depth as many visuo-motor tasks rely on the coordination of sSC and dSC to identify visual information and initiate appropriate behavior. Prey capture is a prime example of an ethologically relevant visuo-motor behavior that relies on the integration of sSC responses to visual input to identify prey and dSC motor output to successfully orient towards, pursue and capture prey^13,14^. In fact, distinct cell types within SC have been identified to contribute differently to prey capture behavior^13^. The cooperation of sSC and dSC can also be seen in escape behavior^15,16^, where it has been shown that sSC identifies looming stimuli^15^ while circuits in dSC initiate increased attentiveness or restlessness associated with escape behavior^17^. Additionally, a recent study explored how SC “sifts” visual information across depth, finding strong differences between responses at different depths within SC, such that the deeper layers of SC are more selective for behaviorally relevant stimuli ^18^. These studies emphasize the need to study sSC and dSC together in order to understand SC’s role in visual behavior.

Furthermore, the role of mouse SC during freely moving head and eye movements has not been well studied. Traditionally, neural responses in SC have been studied using head-fixed or head-rotating experimental paradigms which allow for precise experimental control but give limited insight into active visual processing. Head-fixed neural recordings in sSC demonstrate robust cell type-specific responses to various visual stimuli^3,13^. In dSC, freely moving experiments have found neurons tuned for different directions of head movements around the three axes of rotation^7^ along with tuning to kinetic visual features aligned to movement^19^. However, the range of neural dynamics corresponding to visual and motor representations in SC during free movement has not been directly addressed.

Notably, recent work in mouse primary visual cortex during free movement found that neural responses to gaze-shifting head and eye movements have unique temporal dynamics^20^. These dynamics are visually dependent and correspond with “coarse-to-fine” visual processing, wherein lower spatial frequency (SF) components of the visual scene are processed prior to higher SF components. Importantly, these response dynamics are largely the result of the abrupt change in the visual input that occurs during saccadic movements, rather than encoding the head or eye movements themselves. It is unknown if similar dynamics exist in SC, considering the different roles of collicular and cortical regions in visual processing^4^ and SC’s known involvement in the generation of head and eye movements ^8^.

These previous findings highlight the importance of studying vision in unrestrained conditions allowing for head and eye movements. While there is extensive research studying the visual responses in sSC and motor responses in dSC, it is not yet clear how these regions respond together during naturalistic active vision. We therefore studied neural dynamics across the depth of SC during unrestrained freely initiated head and eye movements.

## Results

### Measuring head and eye movements together with neural activity in SC of freely moving mice

In order to characterize the responses of neurons in mouse SC across depth during head and eye movements, we used chronic electrophysiology and a head-mounted camera system we previously employed to study V1 responses^20,21^. Briefly, this system enables measurement of pupil position with a camera aimed at the right eye, the animal’s field of view with a forward-facing camera, head rotation with an inertial measurement unit (IMU), and single-unit electrophysiology with a chronically implanted 128-channel silicon electrode positioned in SC (Figure 1A). Our set-up allowed for sequential head-fixed and freely moving recordings, enabling us to measure responses in the same units to controlled visual stimuli during head-fixation, and during active sensing in free movement (Figure 1B). To determine electrode depth, in addition to post-hoc histology (Figure 1C), we presented contrast-reversing checkerboard stimuli to measure the local field potential (LFP). It has been previously demonstrated that this stimulus elicits a maximal response in the superficial layer of the SC^22^. We therefore determined the depth of each electrode channel by referencing to the center of the superficial layer, based on the channel with maximal LFP deflection (Figure 1D, E).

**Figure 1.**
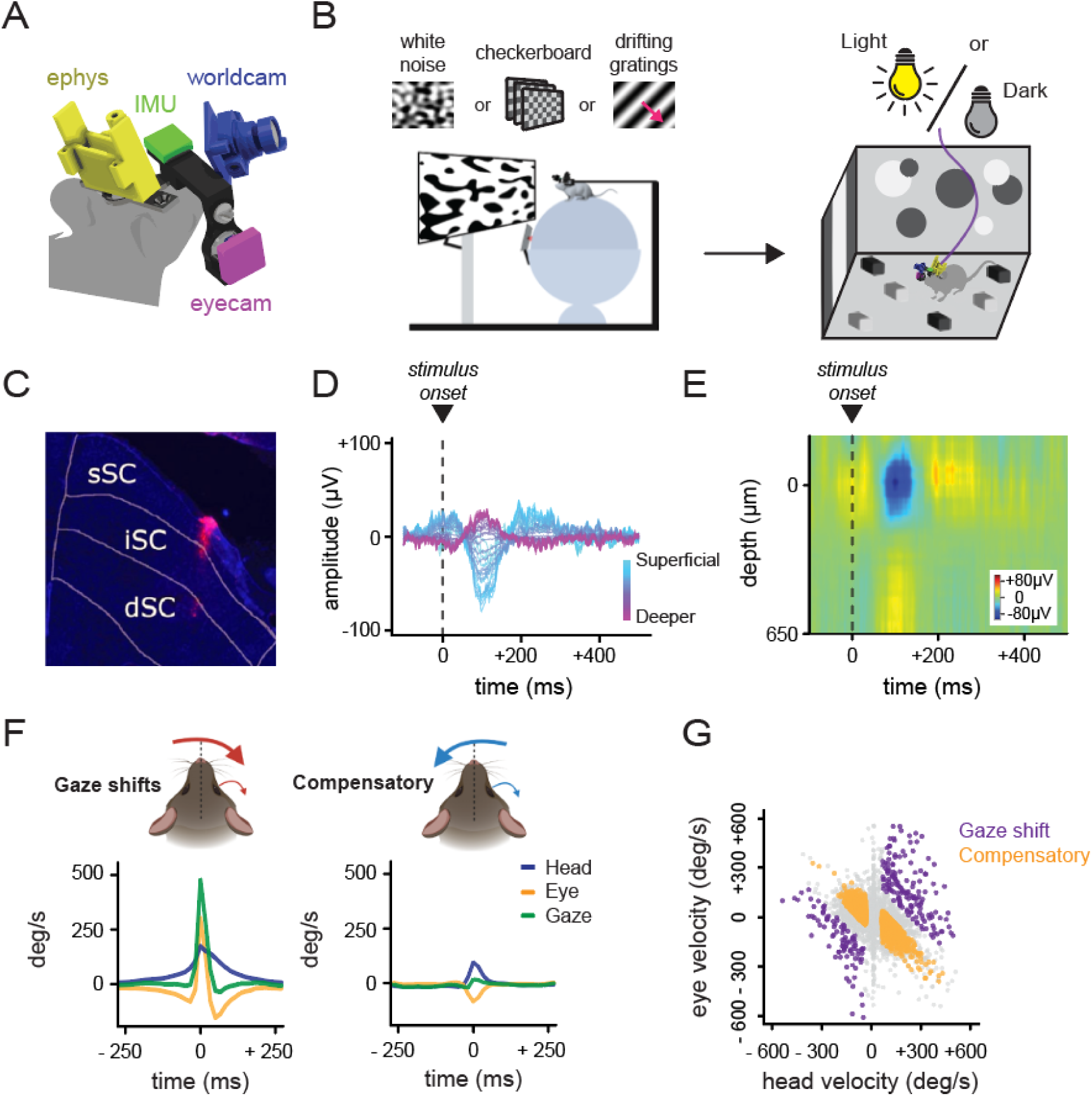
Experimental configuration. (A) Schematic of head-mounted recording system including 128 channel silicon electrode implanted in SC, eye camera, world camera, and inertial measurement unit (IMU). (B) Experimental paradigm where neural responses are recorded during head-fixed set-up that consists of white noise, checkerboard, and drifting grating sessions, immediately followed by freely moving recordings, light sessions (n = 11 animals) and dark sessions (n = 6 animals) (C) Histology of electrode tract (magenta) going through all layers of SC. (D) Local field potential (LFP) waveforms for example electrode shank showing amplitude responses for each channel after checkerboard stimulus presentation (shown by dashed line) with deeper cells in purple and more superficial cells in cyan. Depth was calculated based on maximum amplitude deflection described in methods and modeled after Ito et al., 2017^22^. (E) Heatmap of LFP waveforms for each channel sorted by depth. Dashed line indicates the onset of the reverse checkerboard stimulus. (F) Schematic of the two primary types of head/eye movements, along with the averaged trace for head, eye, and gaze (the summation of eye and head movements). (G) Scatter plot of eye and head velocities, from the example recording used in A, showing compensatory, gaze-shift, and intermediate movements (intermediate shown in grey).

Mice sample the visual scene in a ‘saccade-and-fixate’ pattern, found across the animal kingdom, in which coordinated head and eye movements select and stabilize input to the retina^23^. During head movement, the mouse’s eyes perform one of two actions: move in the same direction as the head to shift its gaze, or move in the opposite direction of the head to stabilize gaze; we refer to these movements as gaze-shifting or compensatory, respectively^24,25^. During free movement these behaviors occur in a continuous pattern that represents rapid gaze-shifting head/eye movements (saccade) and ongoing stabilizing head/eye movements (fixate). In this work, we examined only horizontal gaze-shifting movements during free movement since vertical eye movements in mice are largely compensatory^24–27^.

Consistent with previous findings during free movement^20,24,25^, we observed many large amplitude horizontal head movements that were accompanied by horizontal eye movements. We classified these movements as either gaze-shifting or compensatory by calculating the gaze velocity (i.e. sum of head and eye movement). Gaze-shifting movements result when the animal moves its head and eyes together in the same direction to shift their gaze (Figure 1F; left), whereas compensatory movements result when the animal moves its head and eyes in equal but opposite directions to stabilize gaze (Figure 1F; right). This categorization can be easily visualized by plotting eye vs head velocity, where compensatory movements align along the diagonal, representing equal but opposite head and eye movements, whereas gaze-shifting movements fall beyond the diagonal (Figure 1G). We define gaze-shifting and compensatory movements as previously outlined^20^; briefly, timepoints where gaze velocity is >240 deg s^-1^ are gaze-shifting movements and timepoints where gaze velocity is <120 deg s^-1^ are compensatory movements. Movements that had gaze velocity values between 120 deg s^-1^ and 240 deg s^-1^ were excluded from analysis to avoid ambiguity in classification.

### Temporal dynamics of gaze shift responses in SC

From the silicon probe recordings, we extracted single-unit neural activity across the depth of SC in freely moving mice around the time of gaze-shifting and compensatory head and eye movements. In our population of 542 neurons from 11 mice, we find a majority of cells (57.9%; 314/542) respond to these gaze-shifting movements, which produce abrupt changes in retinal input, whereas responses to compensatory movements, which stabilize retinal input, are more rare (22.0%; 119/542). Figure 2A,B shows example responses of neurons in sSC and dSC to gaze-shift movements. The sSC neurons responded with a positive or negative change in firing rate immediately following the gaze shift, and their response was generally similar for both directions of gaze shifts. On the other hand, the dSC neurons began responding before the gaze shift with a more extended duration, with a strong positive response to one direction of gaze shift and a negative response to the other direction. This response type difference can also be seen in individual continuous recordings when comparing rasters of neural responses at the population level during free movement; sSC neurons respond in a unique temporal dynamic at the onset of a gaze-shift (Figure 2C, bottom) while in contrast dSC neurons respond predominantly in a synchronous burst-like fashion that initiates prior to the gaze-shift and persists after (Figure 2D, bottom).

**Figure 2.**
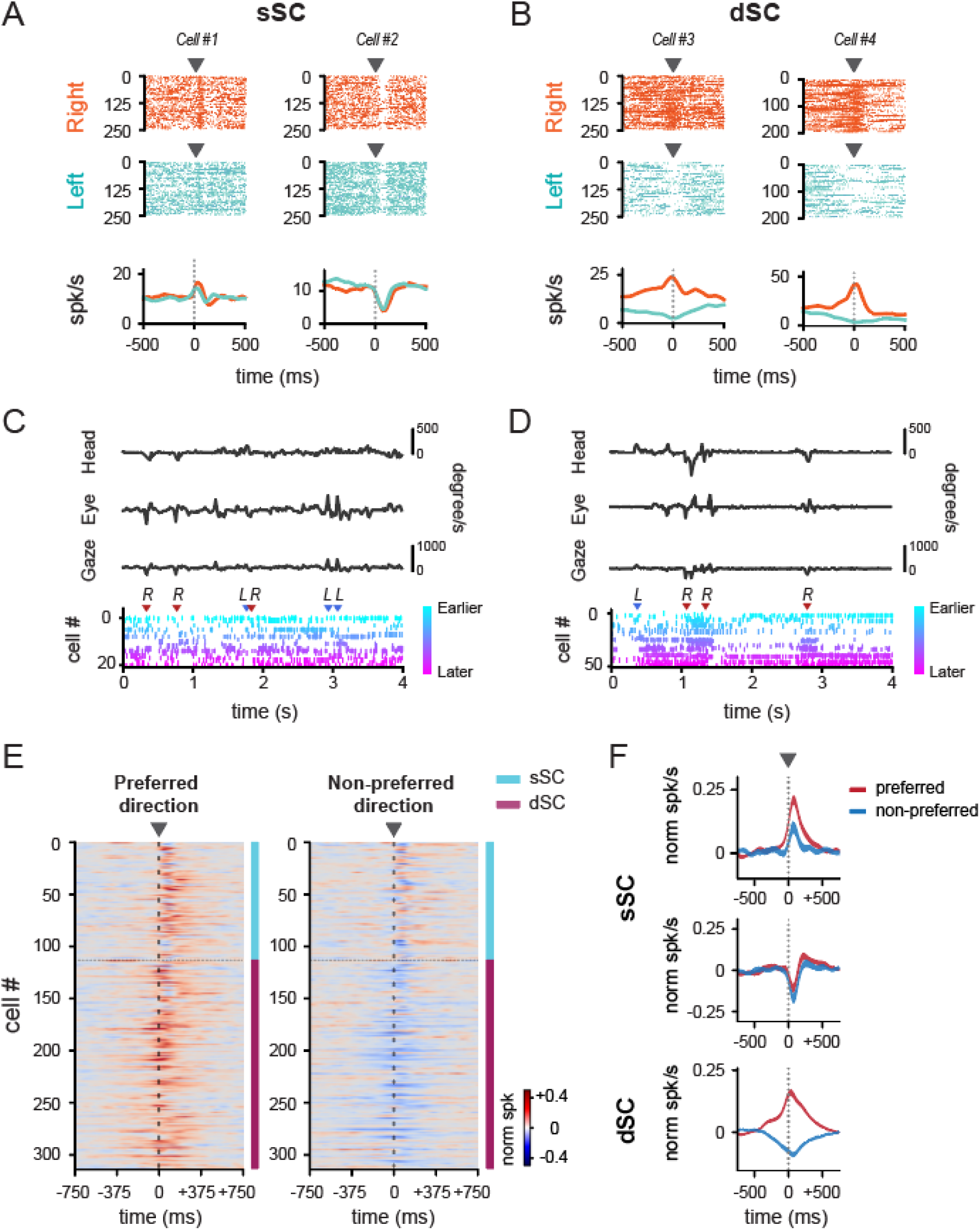
Temporal dynamics of gaze shift responses in SC. (A) Example spike rasters showing two different response types: positive (*left*) and negative (*right*) in sSC for rightward gaze shift movements (*top*), leftward gaze shift movements (*middle*) and corresponding PETHs (*bottom*), rightward gaze shift movement (*orange*) and leftward gaze shift movement (*cyan*). Arrowheads indicate onset of gaze shift. (B) Same as A but for dSC motor responsive example neurons. (C) Example traces showing head movement, eye movement, and gaze (sum of head and eye movements). Bottom raster shows responses from neurons in sSC during this experiment. Gradient colors indicate latency to gaze shift onset. L and R arrows indicate time of gaze shift, and arrow color indicates which direction gaze shift was, red for right, blue for left. (D) Same as E but for example dSC experiment. (E) Temporal sequence of preferred gaze shift responses (*left*) and non-preferred gaze shift responses (*right*) across depth, organized by depth, with cell 0 being the most superficial. (n = 11 mice, 314 cells) (F) Mean PETHs response for gaze-responsive sSC positive cells (*top*; n = 81 cells), gaze-responsive sSC negative cells (*middle*; n = 34 cells), and dSC gaze-responsive cells (*bottom*; n = 199 cells). Red line shows the preferred gaze shift response. Blue line shows non-preferred gaze shift direction response.

The temporal dynamic resulting from gaze shifts is further distinguished between sSC and dSC when looking at population-level temporal sequences sorted by depth from most superficial sSC to deepest dSC, as shown in Figure 2E. The most superficial cells respond quickly after the onset of the gaze shift with either a positive or negative change in firing rate, whereas cells in the dSC begin to respond around 500 ms prior to gaze shift onset, and persist for 500 ms after. Neurons in sSC generally respond similarly to preferred and non-preferred gaze shift directions whereas neurons in dSC show clear direction preference with positive responses to preferred gaze shifts and negative responses to non-preferred gaze shifts (Figure 2E). We further quantified these differences based on the mean peri-event time histograms (PETHs) across the population in response to gaze shifts in sSC and dSC. In sSC, neurons have either positive or negative responses consistent across the two directions of gaze shift, while neurons in dSC show opposite responses to the two directions of gaze shift (Figure 2F). Additionally, while sSC neurons are not responsive to compensatory movements, dSC neurons do respond to compensatory movements (Figure S1B-C). However, we see smaller amplitude responses during compensatory movements as compared to gaze-shifting movements (Figure S1D). This difference is consistent with the fact that compensatory movements on average have lower head amplitude than gaze-shifting movements (Figure S1E) (*p* < 0.0001; Wilcoxon rank-sum test) (gaze-shift: 373.42 deg s^-1^, compensatory: 285.11 deg s^-1^).

### dSC gaze-shift responses persist in the dark, but sSC responses do not

The above results show that sSC neurons respond following gaze shift onset, largely independent of the direction of gaze shift movement. On the other hand, dSC neurons begin responding prior to gaze shift onset, and they encode the direction of movement. Together, these results are consistent with superficial neurons responding to the change in visual input that occurs during gaze shifting movements and deep neurons encoding the motor aspect of the movements themselves. To directly test the role of visual input in these dynamics, we recorded gaze shift responses from the same neurons during free movement in both light and dark conditions. Gaze shift and compensatory head/eye movements occur during dark sessions similarly to those in the light (Figure S2A). However, in darkness, the neural response following a gaze-shift changes drastically for sSC cells whereas the response of dSC cells remains nearly identical (Figure 3A, B). Specifically, both positive and negative sSC cell responses are largely eliminated in darkness, while dSC responses are unaffected (Figure 3B). These differences are also evident in the mean PETH gaze shift responses in light and dark (Figure 3C). Thus, during active vision sSC and dSC respond differently based on visual input, with sSC responding largely to the saccadic gaze shifts in a visually dependent manner, and dSC responding to both gaze-shifting and compensatory movements even in the absence of visual input, consistent with encoding the motor component itself.

**Figure 3.**
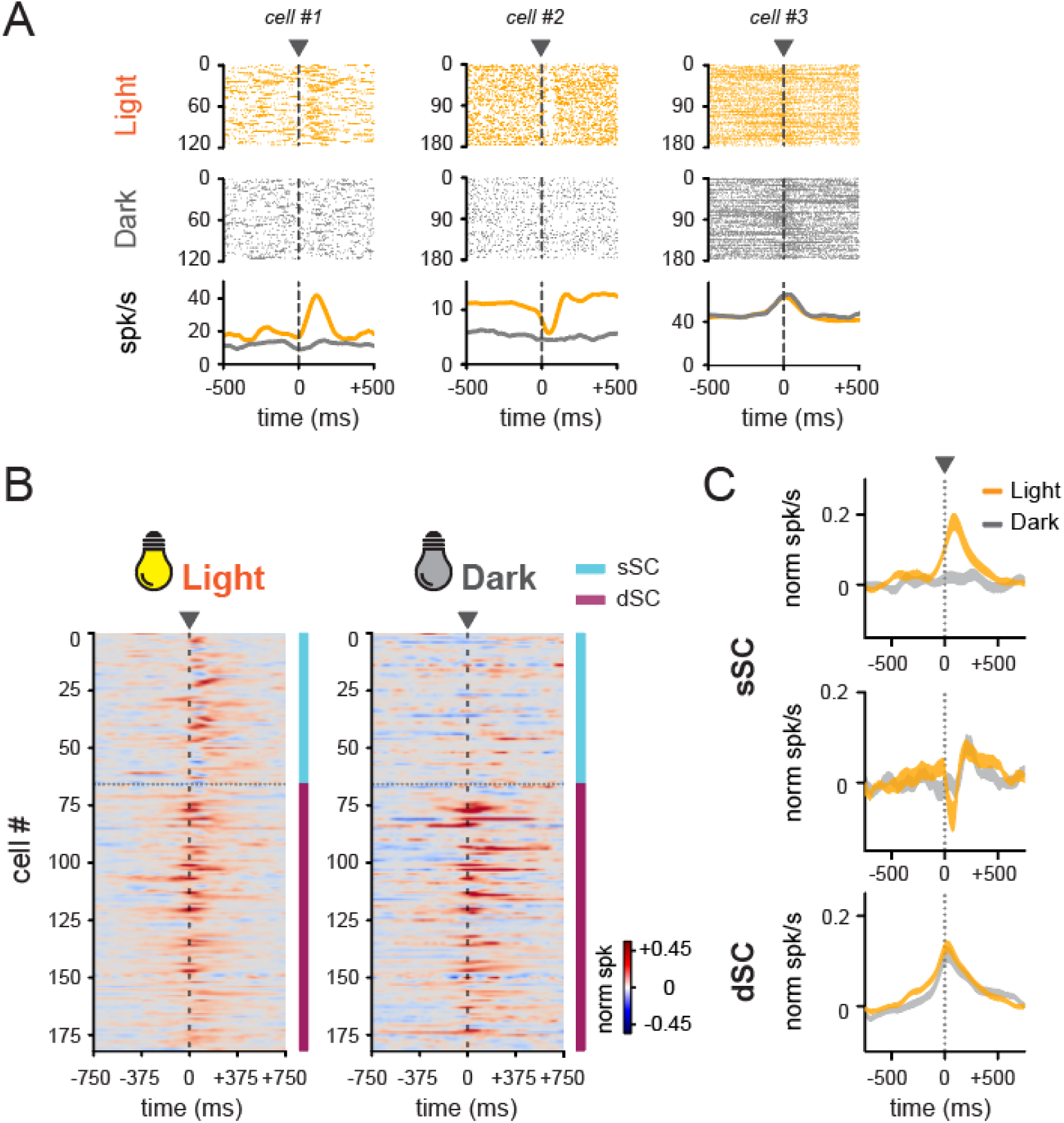
sSC gaze shift responses are visually driven, dSC are not. (A) Example spike rasters from light (*top*) and dark (*middle*) conditions, and their corresponding PETHs (*bottom*). Arrowheads indicate onset of gaze shift. (B) Temporal sequence of gaze shift responses during light (*left*) and dark (*right*) recording sessions plotted by depth (n = 6 animals, 182 cells). (C) Mean PETH of sSC early positive (*top;* n = 40 cells), negative (*middle;* n = 25 cells), and dSC (*bottom;* n = 117 cells) cells in light and dark conditions.

### sSC neurons are tuned to visual contrast while dSC neurons are tuned to head movement

We next sought to further investigate the relative responsiveness of SC neurons to visual input versus head movement itself. To measure visual responsiveness, we recorded from the same units during head-fixation, while presenting visual stimuli on a computer monitor. We used a band-limited white noise stimulus, which was contrast modulated such that it progresses from grey screen up to full contrast and back down to zero contrast over a period of 10 seconds^28^. This stimulus allowed us to measure the contrast response function of each neuron based on the temporally varying contrast level. We observed a clear demarcation in visual responsiveness between sSC and dSC. Figure 4A shows contrast response functions measured from white noise for the same example sSC and dSC units as in Figure 2A,B. The positive and negative sSC cells have strong contrast responses, whereas no distinct response to visual contrast is seen in the dSC neurons (Figure 4A).

**Figure 4.**
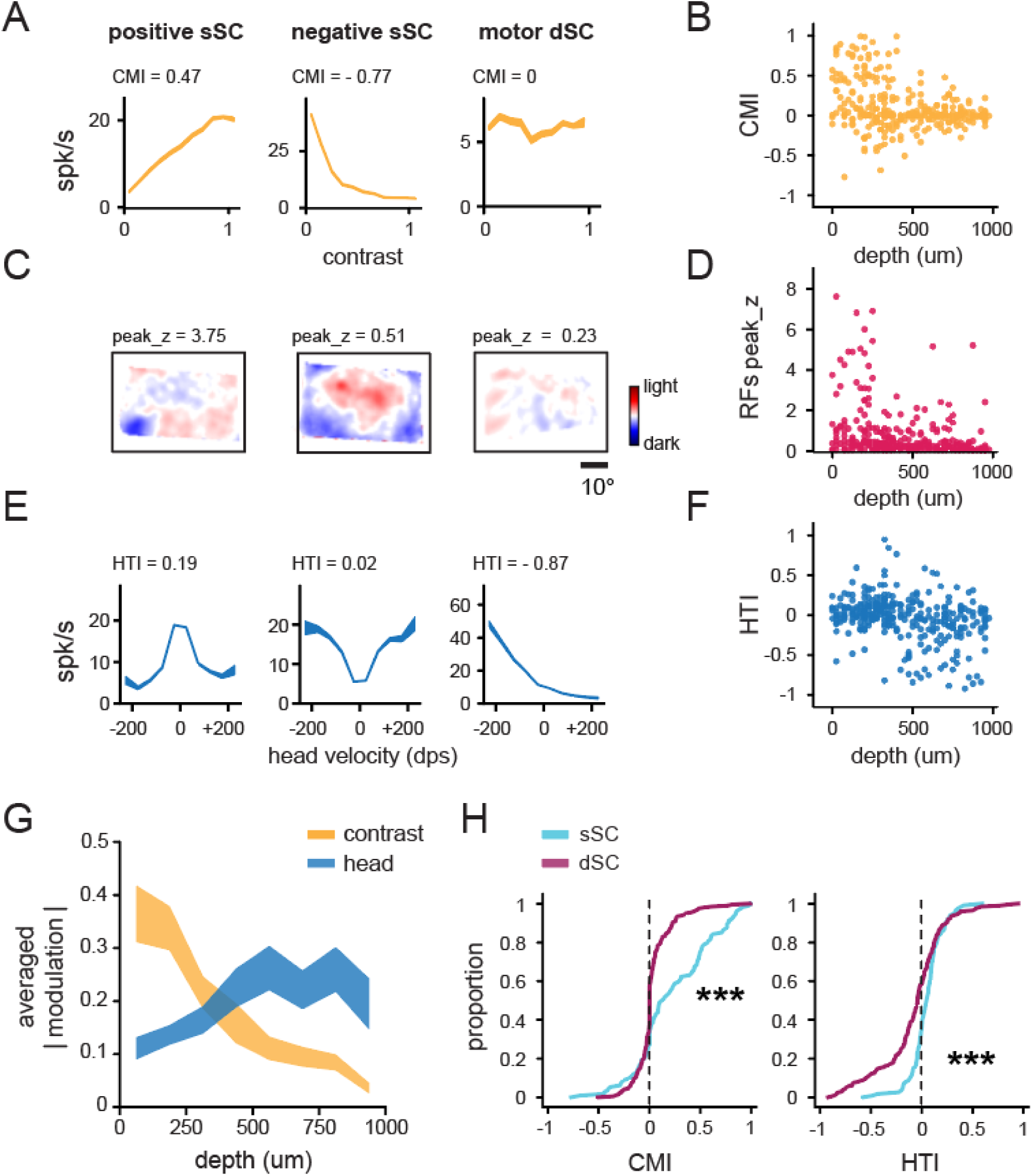
sSC is tuned to visual contrast, dSC is tuned for head-movement. (A) Contrast response functions for each response type, positive sSC, negative sSC, motor dSC, calculated from response to presentation of contrast variable white noise stimulus. (B) Contrast modulation index (CMI) is calculated using cells’ response to contrast-modulated white noise stimulus ((R_CMax_ - R_C0_)/(R_CMax_ - R_C0_)). (C) Receptive field (RF) maps for each of the example units in A. RF was calculated from spike triggered average responses to white noise stimulus. Scale bar for RFs represents 10°. (D) Z-score scatter calculated from white noise spike triggered average. (E) Head velocity tuning curves from example SC neurons. (F) Head-movement tuning index (HTI) was calculated for each cell ((R_+360°_ - R_-360°_)/(R_+360°_ + R_-360°_)). (G) Average modulation (absolute value) of SC neurons across depth to CMI and HTI). (H) Cumulative distribution of CMI and HTI between sSC and dSC (n = 11 mice, 314 cells). *** < 0.001.

We further quantified contrast responses of the population across SC depth by calculating a contrast modulation index (CMI) for each gaze-shift responsive cell based on the response to maximum contrast R_max_ and to zero contrast R_0_ as ((R_CMax_ - R_C0_)/(R_CMax_ + R_C0_)). Again a clear distinction in the degree of contrast modulation emerges across SC depth. We see that cells in sSC have varying degrees of contrast modulation ranging from 1 to -1, whereas dSC do not appear to be modulated by contrast (Figure 4B).

The white noise stimulus also allowed us to identify a neuron’s receptive field based on the spike-triggered average. Example neurons in sSC show robust receptive fields whereas the dSC neuron lacks a defined receptive field (Figure 4C). These RF trends persist at a population level (Figure 4D). Visual contrast and spike triggered average responses from head-fixation suggest that gaze-shift responsive neurons in sSC are responsive to white noise visual stimuli whereas dSC cells are not. Together with the fact that sSC responses are eliminated in the dark while dSC responses persist further suggests that sSC gaze shift responses are driven by the visual input while dSC neurons are driven by the head movement themselves. However, it should be noted that white noise may not be an effective stimulus for all neurons, and the lack of dSC responses could also indicate a different visual feature selectivity.

We next examined the degree to which sSC and dSC neurons encode head movement by calculating a head velocity tuning curve for each cell in SC. Figure 4E shows head velocity tuning curves for two individual sSC neurons and one dSC neuron. While sSC neurons respond with either activation or suppression to head movements in both directions, the dSC unit shows strong head direction preference in one direction. We further quantified head velocity tuning by calculating a head-movement tuning index (HTI) for each cell based on its firing rate at +360 deg/sec and -360 deg/sec as ((R_+360_ - R_-360_)/(R_+360_ + R_-360_)) (Figure 4F). Across the population, only a small fraction of cells in the sSC appeared to be tuned to specific directions of head movement (19.1%; 22/115), whereas a large proportion of dSC cells are modulated by a specific direction of head movement (cells whose firing rates are modulated>50% in one direction: 36.7%, 73/199; Figure 4F). Specifically, among these head movement tuned cells, the majority preferred a rightward turn (i.e., negative angular velocity), corresponding to the contralateral side of the probe implant (rightward cells: 68.5%, 50/73; leftward cells: 31.5%, 23/73; Figure 4F). Furthermore, we also found that this head movement tuning was preserved for dSC cells in darkness (Z = 1874, *p* = 0.727; Wilcoxon signed-rank test) (Figure S2B).

As a final comparison between responses to visual stimuli and head movement, we calculated the mean CMI and HTI across depth in the SC (Figure 4G). A clear distinction between visual and head movement modulation across depth was evident. sSC cells are more strongly modulated by visual contrast than dSC cells (*p* < 0.0001; two-sample Kolmogorov-Smirnov test), while dSC cells are strongly modulated by head movement in one direction, whereas sSC cells are not (*p* < 0.0001; two-sample Kolmogorov-Smirnov test) (Figure 4H).

### sSC neurons have unique gaze shift initiated temporal dynamics when compared to V1

Based on the fact that sSC unit responses were dependent on visual input, we examined their temporal dynamics by sorting units based on the latency of their peak positive response to gaze shifts, as in Parker et al., 2023 for V1. This revealed a clear temporal sequence in sSC in the mean response to gaze shifts (Figure 5A). We confirmed that the resulting temporal sequence was not an artifact of sorting through cross-validation (Fig. S3; Pearson correlation coefficient, r = 0.67, *p* < 0.0001). Over 50% of the cells in the sSC had a peak latency within 100 ms, demonstrating that their response occurs rapidly following gaze shift events (Figure 5B). This temporal sequence response in the sSC can also be visualized in single-trial spike rasters (Figure 2E).

**Figure 5.**
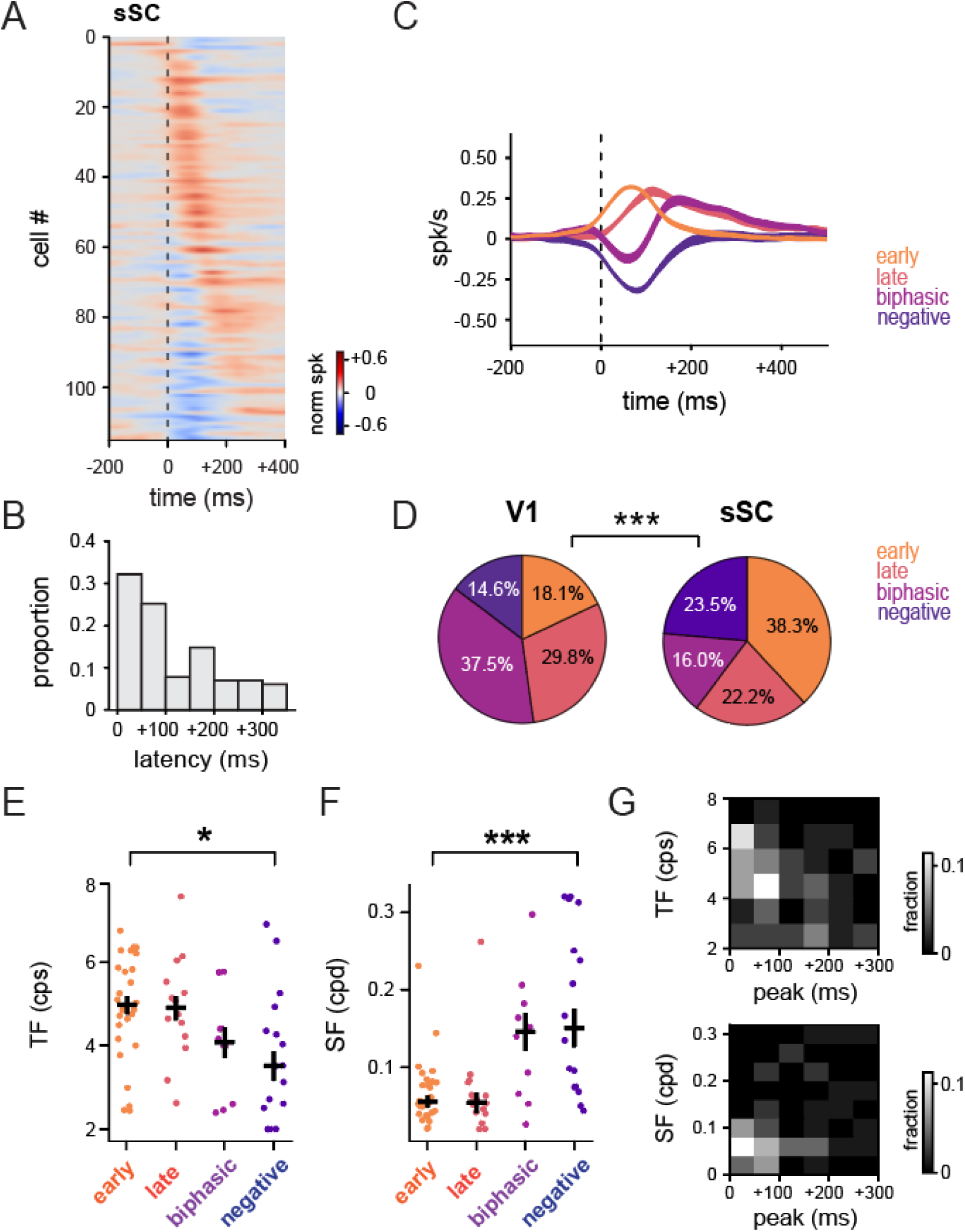
Coarse-to-fine processing in sSC. (A) Temporal sequence of gaze shift responses for all gaze-responsive cells in sSC, plotted by peak latencies (n = 11 animals, 115 cells). (B) Distribution of the latencies of positive peaks in gaze-shift PETHs for each unit. (C) Mean PETHs of sSC response types clustered using the same metrics as in Parker et al., 2023. (D) Proportion of cells in each response type between V1 and sSC showing significant difference in proportion of cells in each response group. (Chi-square test X^2^_(1)_= 42.12, *p* < 0.001). (E-F) Weighted TF (left) and SF (right) preferences for each cluster, calculated as a weighted mean of responses for two presented TFs and four presented SFs. (G) Distribution of TF (*top*) and SF (*bottom*) across the peak latencies of the clustered cells.

In order to directly compare sSC to V1, we applied the same clustering as previously used to segregate V1 neurons^20^. Briefly, we performed *k*-means clustering of units based on principal component analysis (PCA) of the normalized PETHs for gaze shifts in the preferred direction, using the same principal components and *k*-means model from V1. Clustering allows us to categorize the heterogeneous population dynamics of sSC, similarly to V1, however, it should be noted that we are subdividing a continuous population. A value of *k* = 5 clusters was implemented to allow for direct comparison to V1 gaze shift dynamics in Parker et al., 2023. This resulted in four responsive clusters, early positive (24.6%), late positive (14.3%), biphasic (10.3%), and negative (15.1%) responses to gaze shifts, as well as a large unresponsive fifth cluster (35.7%) (Figure 5C). It should be pointed out that while all four response types appear in sSC this is a direct consequence of using the same *k*-means clustering model as used for V1 experiments ^20^. However, clustering within SC revealed that the response types were not strikingly different from those obtained using the V1 model (Figure S4). When directly comparing the distribution of these response types within sSC, however, the population is significantly different compared to V1. While we found no difference in the fraction of gaze shift responsive cells between V1 (63.8%) and sSC (64.3%) (Chi-square test X^2^_(1)_= 0.04, *p* = 0.83), in the sSC population there are significantly more early positive responsive cells and fewer biphasic responsive cells (Chi-square test X^2^_(1)_= 42.12, *p* < 0.001; Figure 5D). Together, these results suggest that sSC and V1 respond to gaze shifts with different temporal dynamics.

### Coarse-to-fine processing around gaze shifts in sSC

We sought to further investigate the visual tuning properties of sSC gaze shift responsive cells. Specifically, we were interested to see if cells in sSC had a coarse-to-fine relationship between their visual tuning responses and temporal responses to gaze shifting head/eye movements, similar to that observed in V1^20^. During head-fixation we presented drifting sinusoidal gratings at three spatial frequencies (0.02, 0.08, 0.32 cycles per degree (cpd)) and two temporal frequencies (2, 8 cycles per second, cps). We first compared temporal frequency (TF) preferences for each response type in sSC, and found a significant difference in TF between clusters (Figure 5E; X^2^_(3)_ = 10.025, *p* = 0.018, Kruskal-Wallis test, n = 11, n = 71 cells). Across all clusters, early cells had significantly higher TF preference compared to biphasic and negative cells (early vs. biphasic: Z = 2.254, *p* = 0.024; early vs. negative: Z = 2.391, *p* = 0.017; Wilcoxon rank-sum test), but had similar TF preferences to late cells (Z = 0.374, *p* = 0.709; Wilcoxon rank-sum test). Additionally, there were no differences between biphasic and negative cells (Z = 0.598, *p* = 0.527; Wilcoxon rank-sum test). This suggests that there is a temporal frequency preference difference across sSC cell types, but unlike in V1, they do not follow a gradually increasing pattern, but rather a more discrete pattern (i.e., low TF: early & late; higher TF: biphasic & negative).

We next compared SF preferences in sSC for different gaze shift response types. We found a significant difference in SF preferences across each k-means clustered response type in sSC (χ2_(3)_ = 19.540, p < 0.001; Figure 5F). Furthermore, this trend was consistent between sSC positive and sSC negative cells, which is independent of the *k*-means clustering method. Positive cells exhibited higher TF than negative cells (Z: 3.278, *p* = 0.001; Wilcoxon rank-sum test; Figure S5A), while having lower SF than negative cells (Z = -3.217, *p* = 0.0013; Wilcoxon rank-sum test; Figure S5B).

Unlike the broad distribution of SF and TF found in V1, we mainly observed high TF/low SF with weak low TF/high SF at the population level in sSC (Figure 5G). This suggests that the sSC might be more involved in fast and coarse visual information processing. Another notable difference was a significant population of cells with biphasic/negative (i.e., sSC negative cells) responses to gaze shift, which were tuned to to high TF/high SF, rather than the inverse relationship between these associated with coarse-to-fine (Figure S6). Similar response properties of units have been described for stellate cells in the SC^3^, suggesting that these cells may correspond to a population that does not follow the typical coarse-to-fine processing. Together, these results suggest that sSC processes visual input following gaze-shifts differently than V1.

### Head movements predominate over eye movements in the mouse dSC

Previous studies have demonstrated that saccade-related neurons in the SC of non-human primates exhibit a burst of spikes tightly locked to the onset of eye movement^9,29,30^. To further dissociate the impact of head movement from eye movement in the SC of freely moving mice, we detected burst spike events throughout each freely moving recording session and then averaged head or eye movements based on the onset of the bursts^7^. This approach allowed us to examine each cell’s tuning properties to head or eye movements, dissociating their interrelated patterns to determine which movement—head or eye—was more important for each cell.

First, we examined the fraction of spikes that participated in bursts. We defined bursts as periods during which a neuron fired at least three spikes, with inter-spike intervals not exceeding 20 ms and a minimum burst duration of 20 ms. This fraction was calculated by dividing the spikes during the bursts by the total number of spikes occurring during any head movements (>1 deg s⁻¹). For this analysis, we included cells with at least 30 bursts throughout the session (67.0% of sSC cells; 77/115, 53.8% of dSC cells; 107/199).

Next, we calculated the head and eye movement patterns for each cell around the bursts, from -0.5 s to 1 s, and each cell’s traces of burst-triggered average (BTA) head and eye movements were visualized (Figure 6A, 6B, and 6C). At the population level, for the sSC positive responsive cells, the peak amplitude of head and eye movements preceded the bursts, and their peaks were comparable (Figure 6D). In contrast, in sSC negative cells, the effects of both head and eye movements were minimal during the bursts (Figure 6E). Lastly, dSC cells exhibited more pronounced head movements at the onset of bursts compared to eye movements, suggesting they are more likely to be encoding head movements rather than eye movements (Figure 6F).

**Figure 6.**
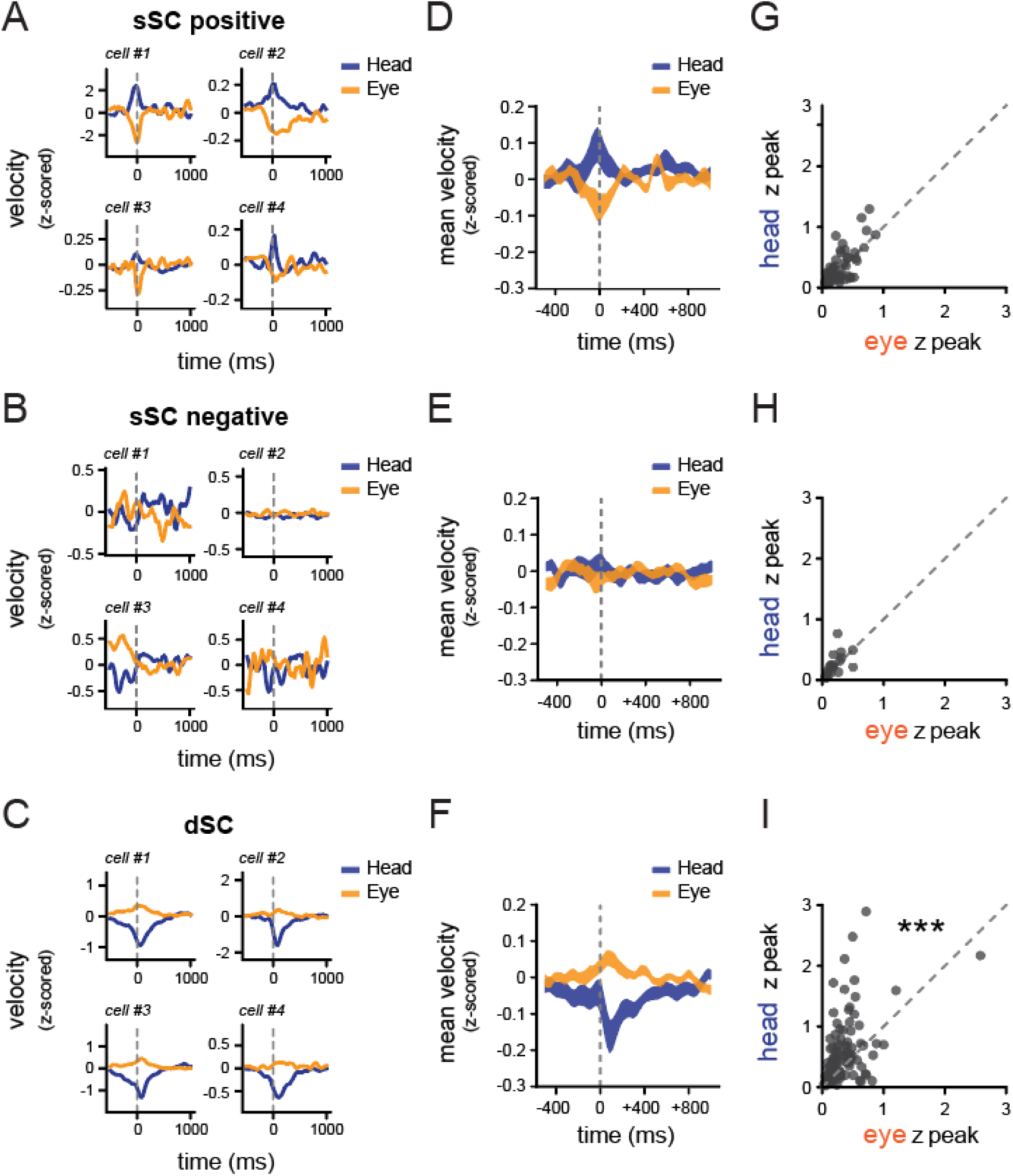
Head movements have a greater influence than eye movements in driving dSC cells. (A-C) Representative burst-triggered averages (BTA) of head (*orange*) or eye (*navy*) movements, aligned to the onset of the burst train of spikes from (A) sSC positive (n = 59 cells), (B) sSC negative (n = 18 cells), and dSC (n = 107 cells). (D-F) Average velocity of head or eye movements from the (D) sSC positive, (E) sSC negative, and (F) dSC populations. Positive values indicate leftward movements, while negative values are rightward. (G-I) Distributions of the absolute peak values of BTA traces from eye vs. head from the (G) sSC positive, (H) sSC negative, and (I) dSC cells.

Lastly, to quantify whether each cell is more tuned to head, eye, or equally tuned to both head and eye movements, we determined their movement preferences by identifying the peak values from each cell’s BTA traces of head and eye movements. We reasoned that a greater peak value from the BTA traces of head or eye movements would play a more dominant role in driving SC units. We detected the absolute peak values of head and eye movements between 0 and +250 ms relative to the bursts. As a result, we observed that the majority of cells were equally driven by both head and eye movements in sSC (45.5%; 35/77), although there was a tendency for both sSC positive cell and sSC negative cells to be slightly more driven by head movements than by eye movements (Figure 6G and Figure 6H) (sSC positive: z = 634.0, *p* = 0.058; sSC negative: z = 46.0, *p* = 0.085 ;Wilcoxon’s signed-rank test). In contrast, we observed a significant skew toward head movements over eye movements in deeper SC cells (Figure 6I) (z = 1397.0, *p* < 0.0001; Wilcoxon’s signed-rank test). While we also observed cells driven by eye movements rather than head movements in both the sSC and dSC, their numbers were relatively small (sSC: 18.2%; 14/77; dSC: 15.6%; 17/107) compared to the head-driven cells (sSC: 36.4%; 28/77; dSC: 50.5%; 54/107). Furthermore, we found no differences in the proportion of eye-driven cells between sSC and dSC (Chi-square test X^2^_(1)_ = 0.39, *p* = 0.53), while there was a significant increase in head-driven cells in dSC compared to sSC (Chi-square test X^2^_(1)_ = 7.96, *p* = 0.0048). These results suggest that head movements are predominantly encoded in the dSC of freely moving mice within our recording environment.

## Discussion

We recorded the activity of neurons across the depth of SC in freely moving mice to investigate how active vision, in terms of head and eye movements, influences neural activity in the superior colliculus. We found distinct gaze-shifting responses for sSC and dSC, with sSC responding to the visual aspect of gaze shifts and dSC responding to the motor aspect of gaze shifts. Specifically, in sSC we find two main response types, positive and negative, to gaze-shifting head/eye movements; however, they are not responsive to compensatory head/eye movements. Additionally, the gaze shift responses of sSC neurons were eliminated in darkness, and sSC neurons have a strong response to white noise stimuli presented during head-fixation. These findings suggest that sSC responses to gaze-shifting head/eye movements likely result from the change in visual input during these saccadic movements as opposed to motor efference signals of the combined head/eye movement. On the other hand, we found that neurons in dSC are tuned to the direction of head movement, in both light and dark, and have minimal response to white noise stimuli, suggesting that they are encoding the motor component of the gaze shift. These findings are consistent with the established organization of SC, with sSC being visual, and dSC being motor-related^8,9,19,31^, and demonstrate this correspondence during active vision where sensory and motor signals are intertwined.

In primates, dSC is involved in the generation of saccadic eye movements as well as head movements^9,32^ but here we find dSC neurons in mice primarily respond around head movements, with relatively few cells being specifically correlated to eye movements. This difference may be due to key variations in orienting behaviors between species, as in mice head and eye movements are often coupled and they orient primarily with head movements^24,33^. This coupling is reflected in recently described neural circuitry that coordinates head and eye movement in SC output^12^. Furthermore, the finding that most mouse SC neurons do not primarily encode eye saccades is consistent with the notion that many saccadic eye movements in mice are driven by VOR “reset”, as lesion studies in cat and primate suggest that SC does not play a causal role in VOR-coupled saccades^34,35^. However, it remains possible that rapidly appearing or highly salient stimuli could elicit a different form of gaze shifts, initiated by eye movements as previously shown in primates^36^, which could be driven by SC.

We also compared sSC gaze shift responses to previously established coarse-to-fine gaze shift dynamics in V1^20^. Coarse-to-fine visual processing postulates that the visual system responds first to coarse aspects of the visual scene before responding to fine details^37^. We found that sSC neurons that responded earliest following the onset of gaze shift preferred low SF, and neurons responding latest to gaze shift onset preferred higher SF. These results correspond with a weak form of the coarse-to-fine visual processing we observed in V1. sSC spatial frequency preferences clearly differ when compared to V1, which suggests that sSC and V1 process the visual input following a gaze shift differently. While it is not clear what exactly these SF preference differences represent in terms of overall visual processing, one possible explanation is that sSC and V1 are primed to respond differently to the gaze shift induced abrupt change in visual input. For instance, sSC may be tuned to respond preferentially to the coarse, low spatial frequency, features of the new visual input to detect salient stimuli, whereas V1 may be more involved with detailed interpretation of the new visual input, through coarse-to-fine processing. This work sets a foundation for future studies exploring circuit dynamics within SC during unrestrained active vision. Previous studies have highlighted the role of specific cell types in SC during head movements^38^ and prey capture^13^, but recent studies have elucidated a range of identified cell types in SC^3,39,40^ that can be related to visual and motor aspects of active vision. Also, while here we measured SC gaze shift responses during spontaneous exploration, it remains to be determined whether these response dynamics are differently modulated during goal-directed behaviors. Ultimately, this work highlights the importance of studying visual processing in unrestrained conditions to reveal the complex dynamics of visual processing, while also providing a basis for future studies to investigate gaze shift dynamics in SC during many behaviors at the cellular and circuit level.

## Author Contributions

SLS and CMN conceived the project. CMN supervised all aspects of the project. SLS led mouse experiments. KJ contributed to histology and mouse experiments. JS led data analysis. SLS and DMM contributed to data analysis. SLS, JS, and CMN contributed to writing and editing the manuscript.

## Acknowledgements

We thank Dr. Spencer Smith, Dr. Michael Stryker, Dr. Phil Parker, and members of the Niell Lab for feedback on previous drafts of the manuscript. This work was supported by NIH grants R01NS127305 (CMN), R01NS121919 (CMN) and F31EY034792 (SLS).

## Methods

### Animals

All procedures were conducted in accordance with National Institutes of Health guidelines and were approved by the University of Oregon Institutional Animal Care and Use Committee. 5-10 month-old mice (Mus musculus, C57BL/6J, Jackson Laboratories and bred-in house) were kept on a 12-h light/dark cycle. In total, five male and six female mice were used for this study. Mice were housed with sibling cagemates until the beginning of experiments and then they were singly housed. Humidity was between 40-60% and temperature was 21±1°C. Data collection and analysis were not performed blind due to the condition of the experiments. No animal or data points were excluded from analysis. Data collection was not randomized as there was only one experimental group.

### Surgery and habituation

Mice were initially implanted with a titanium headplate over SC to allow for head-fixation and attachment of head-mounted experimental hardware. After 3 days of recovery a miniature connector (Mill-Max 853-93-100-10-001000) was secured to the skull to allow for repeated, reversible attachment of a camera arm, eye/world cameras, and IMU^20,21,24^. To simulate the weight of the real electrophysiology implant for habituation, a ‘dummy’ electrophysiology drive was glued to the headplate. Animals were handled for several days before surgical procedures, and habituated (∼60 mins) to the head-fixed spherical treadmill and freely moving arena while tethered for several days before experiments.

The electrophysiology implant was performed once animals moved comfortably for an extended period of time in the arena. A craniotomy was performed over monocular SC, and linear silicon probe (128 channels, Diagnostic Biochips P64-10D, P128-6) mounted in a custom three-dimensionally printed drive (Yuta Senzai, UCSF) was lowered into the brain using a stereotax to an approximate tip depth of 1500 μm from the pial surface. The surface of the craniotomy was covered with artificial dura (Dow DOWSIL 3-4680) and the drive was secured to the headplate using light-curable dental acrylic (Unifast LC). A second craniotomy was performed above the left frontal cortex and a reference wire was inserted into the brain. The opening of the craniotomy was covered with a small amount of sterile ophthalmic ointment and the wire was glued in place with UV light cure glue (Loctite AA3972). Animals recovered overnight and experiments began the following day.

### Hardware and recording

The camera arm was oriented approximately 90 deg to the right of the nose and included an eye-facing camera (iSecurity101 1000TVL NTSC, 30 frames per s (fps) interlaced), an infrared light-emitting diode to illuminate the eye (Chanzon, 3-mm diameter, 940-nm wavelength), a wide-angle camera oriented toward the mouse’s point of view (BETAFPV C01, 30 fps interlaced) and an inertial measurement unit acquiring three-axis gyroscope and accelerometer signals (Rosco Technologies; acquired at 30 kHz, downsampled to 300 Hz and interpolated to camera data). Fine-gauge wire (Cooner, 36 AWG, no. CZ1174CLR) connected the IMU to its acquisition box, and each of the cameras to a USB video capture device (Pinnacle Dazzle or StarTech USB3HDCAP). A top-down camera (FLIR Blackfly USB3, 60 fps) recorded the mouse in the arena.

The Diagnostic Biochips electrophysiology headstage (built into the silicon probe package) was connected to an Open Ephys acquisition system via an ultra-thin cable (Intan no. C3216). Electrophysiology data were acquired at 30 kHz and bandpass filtered between 0.01 Hz and 7.5 kHz. We first used the Open Ephys GUI (https://github.com/open-ephys/plugin-GUI) to assess the quality of the electrophysiology data, then recordings were performed in Bonsai20 using custom workflows (https://github.com/nielllab/FreelyMovingEphys). System timestamps were collected for all hardware devices and later used to align data streams through interpolation.

During experiments, animals were first head-fixed on a spherical treadmill to measure visual tuning properties using traditional metrics, then immediately transferred to an arena where they could move freely and explore the visually enriched environment. Recording duration was approximately 40 min head-fixed and 1 hr freely moving. For head-fixed experiments, a 27 inch monitor (BenQ GW2780) was placed approximately 27.5 cm away from the mouse’s right eye, and visual stimuli were presented using Pyschtoolbox-3^41^. Head-fixed stimuli were recorded using the head-mounted world camera. We first presented 15 min of a band-limited Gaussian noise stimulus^42^ (spatial frequency spectrum 0.05 cpd to 0.12 cpd, flat temporal frequency spectrum with a low-pass cutoff at 4 Hz) to confirm SC targeting based on spike-triggered average receptive fields. We next presented flashed sparse noise stimuli consisting of full- and minimum-luminance circular spots on a gray background played for 5 min. Spots were a range of sizes, 2,3,8,16, and 32 degrees in diameter and were presented so that each size made up an equal fraction of the screen, approximately 15% on average. Each stimulus frame was presented for 250 ms immediately followed by the previous frame without ISI^42^. Next, sinusoidal gratings were presented at eight directions of motion for three spatial frequencies (0.02, 0.08, 0.32 cpd) and two temporal frequencies (2, 8 cps) for 10 min, with a 1 sec stimulus duration and 0.5 sec gray ISI with stimulus conditions randomly interwoven (12 presentations per stimulus). Finally, we presented a contrast-reversing square-wave checkerboard stimulus for 5 min with a spatial frequency of 0.04 cpd and temporal frequency of 0.5 Hz.

The arena for free moving experiments was 48-cm long by 37-cm wide by 30-cm high. Three of the arena walls were custom wallpaper covered by acrylic including black and white high- and low-spatial frequency gratings and white noise, the other wall was covered by a 24 inch monitor (BenQ GW2480). The monitor displayed moving sparse noise where full- and minimum-luminance spots were 4, 8, and 16 deg in diameter. Each spot moved in one of eight evenly spaced directions at one of five speeds (10,20,40,80,120 deg s^-1^). Spots appeared on the edge of the screen and moved across until they disappeared on the opposing side. The arena floor was a gray silicone mat (Gartful) and was covered with a dense layer of black and white Lego bricks to provide 3-D visual contrast. To encourage foraging during the recording small pieces of tortilla chips (Juanita’s) were lightly scattered on the arena floor; however, animals were not food or water restricted.

For dark recordings the entire experimental enclosure was sealed in light-blocking material, and all potential light sources within the enclosure and all external light sources were turned off, to ensure complete darkness. Prior to darkness experiments one drop of 2% pilocarpine HCl ophthalmic solution was applied to the animal’s right eye to constrict the pupil and allow for accurate eye tracking as described in Parker et al., 2023. Once the pupil was restricted enough for tracking in the dark (3-5 min) the animal was moved from the spherical treadmill to the dark arena for the recording. The recording lasted approximately 20 min or until the effect of pilocarpine wore off followed immediately by the light recording.

### Data pre-processing

Electrophysiology data was preprocessed following the same methodology as Parker et al., 2023. Raw electrophysiology data from head-fixed and free moving recordings in the same session were concatenated into a single file for spike sorting, allowing us to track single units across the entire experiment. Common-mode noise was removed by subtracting the median across all channels at each timepoint. Spike sorting was performed using Kilosort 2.5 (https://github.com/MouseLand/Kilosort), and single units were isolated and selected using Phy 2.0 (Phy 2.0 beta 5; https://github.com/cortex-lab/phy) based on numerous parameters: contamination (< 10%), firing rate (mean > 0.5 Hz across entire recording), autocorrelogram, and waveform shape. Following spike sorting the data were then split back out into individual recordings for further analysis.

Laminar depth was calculated from the multi-unit local field potential (LFP) of head-fixed neural responses during presentation of flash checkerboard stimulus which generates strong LFP responses in sSC. To identify the surface of sSC we followed methods from Ito et al., 2017. Briefly, we averaged evoked LFP responses to reversal of checkerboard stimulus and chose the time point when LFP has the maximum negative amplitude to represent the top of sSC. Similar analysis is done when looking across depth of SC to identify most superficial responses in a given recording. Additionally, cells were split into sSC or dSC, cells that may be in intermediate SC (iSC) were grouped with dSC for simplicity.

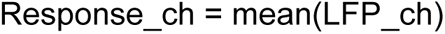

Pupil position was extracted from the eye camera data^24^. Briefly, eye camera data were deinterlaced to achieve 60-fps video, then eight points around the pupil were tracked with DeepLabCut^43^. We then fit an ellipse to these points and computed pupil position by angular rotation. The head-mounted world camera was also deinterlaced to achieve 60-fps video and distortion from the camera lens were corrected with OpenCV. The position of the mouse in the arena was recorded using a top-down camera and tracked using DeepLabCut. Running periods were defined as timepoints when the animal’s neck point had a velocity of > 2cm/s, and stationary states were periods where the animal’s neck point had a velocity of < 2cm/s.

Horizontal head rotation velocity was extracted from the IMU, converted to deg/s and interpolated to eye camera frame timestamps. We defined leftward and rightward directions as the direction of head movement from the animal’s perspective. For example, a movement of the right eye in the nasal direction would be a leftward movement. To select eye/head movement onsets, we first determined if the head velocity was >60 deg/s in the leftward or rightward direction. If a sufficiently large head movement was made, we then separated it into either gaze-shifting or compensatory movements using gaze velocity, where gaze is defined as the sum of horizontal eye and head velocities^24^. If there was a high gaze velocity (>240 deg/s) concurrent with the head movement, the movement was considered gaze-shifting. Those that resulted in low gaze velocity (<120 deg/s) were considered compensatory. Eye/head movements that resulted in intermediate gaze velocities (>120 deg/s and <240 deg/s) were excluded from our analysis to avoid contamination between the two categories of movements. For eye/head movements spanning multiple eye camera frames, only the first time point, representing the onset of the movement, was used to calculate the onset of the PETH. Compensatory movements that occurred 250 ms before or after a gaze-shifting movement were excluded to avoid contamination by gaze shifts. For head-fixed recordings, only high eye velocity (>240 deg/s) was used to identify saccades, since the head was immobilized.

To maintain internal consistency with eye cameras and obviate the need for a separate video stimulus synchronization signal, we determined stimulus onsets in head-fixed recordings directly from the head-mounted cameras. For the reversing checkerboard stimulus that updated every 500 ms, we identified frame transitions from the head-mounted camera video based on *k*-means clustering (*k* = 2) of the video into the two separate contrasts, and selected transitions between the clusters. For the flashed sparse noise stimulus that updated every 250 ms, we determined the timestamps of stimulus onset based on the root mean square pixel-wise change in the image, which showed clear peaks at stimulus transition. For drifting sinusoidal gratings, we determined direction of motion by computing optic flow from the worldcam video, SF based on the mean gradient magnitude from paired Sobel operators and TF from the mean Fourier transform of each pixel over time.

### Analysis of neural responses

All analyses were performed in Python (v.3.8, python.org). Neural responses were calculated as a PETH from electrophysiology spike times using kernel density estimation with a gaussian kernel with bandwidth (standard deviation) of 10 ms, sampled at 1-ms intervals. The PETH included neural activity from −1,000 ms to 1,000 ms around the event onset for all stimuli/events (unless noted otherwise). A modulation index was calculated for PETHs as the peak of the firing rate (*R*) from the event onset at -20 ms until 250 ms after the event, minus the baseline before the event (*Rb*), which was calculated as the mean of *R* from −750 ms to -450 ms and +450ms to +750ms around the event onset (unless noted otherwise).

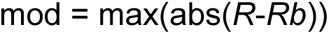

For the drifting gratings stimulus, the PETH was calculated similarly to other stimuli, but from −1,500 ms before until 1,500 ms after stimulus onset to capture the full stimulus and ISI interval.

To normalize PETHs, we subtracted the baseline *Rb* from the PETH (*R*) to give an evoked firing rate, and then divided by the maximum of *R* during a response window (*Rrw*) within a response range of −250 ms before to 250 ms after the event. The denominator includes responses up to 250 ms before the event to ensure that spurious increases in firing rate that are not time-locked to an event were not amplified by the normalization (this was extremely rare and only occurred in unresponsive cells).

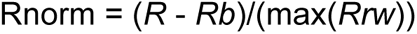

When normalizing the PETH of compensatory eye/head movements, we used that neuron’s *Rrw* from gaze shifts in the preferred left/right direction so that all eye/head responses were normalized relative to the neuron’s best response. Because the head-fixed sparse noise stimulus presentation was of short duration (250 ms), we used the spike rate at the time of the stimulus onset, 0 ms, as the baseline firing rate (*Rb*) for normalization to avoid including responses to previously presented flashed stimuli.

Neurons were considered responsive to a stimulus or eye/head movement onset if they changed their firing rate by at least 10% and 1 spike per second (sp s^−1^) in the 250 ms following the onset of the event. For the gratings stimulus, suppressed-by-contrast cells were not considered responsive, and removed by requiring that gratings PETHs have a peak firing rate >0.5 normalized sp s^−1^ during the 1 s of stimulus presentation.

The latency of a neuron’s peak response was calculated as the timepoint of the maximum firing rate of its normalized PETH in the period between 25 ms to 250 ms after the onset of the event. For temporal sequence plots, normalized PETHs were sorted by the latency of their peak for the preferred direction of gaze-shifting eye/head movements. Peak latency was not calculated for units unresponsive to gaze shifts. Peak latency sorting was cross-validated by randomly assigning gaze-shift events into a train set and a test set, calculating a PETH for each half of the data, and sorting the test set by the peak times of the training set.

SF and TF preferences were based on the evoked firing rate for each stimulus condition, computed as the mean rate from 25 to 1,000 ms following stimulus onset, minus the mean baseline rate in the 500 ms before stimulus onset. The mean SF and TF tuning curves for all cells within gaze-shift clusters were calculated using each cell’s mean evoked firing rate for each SF/TF normalized by the maximum response to its preferred SF/TF.

Each cell’s weighted TF response, *W*_TF_, was determined using the mean evoked responses, *R*, for each of the two presented TFs and at the cell’s optimal SF and preferred orientation.

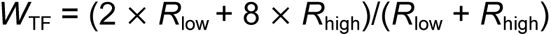

Likewise, weighted SF response was calculated at the cell’s weighted TF response and preferred orientation using the mean evoked responses for each of the three presented SFs.

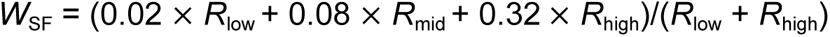

### Response clusters

To compare visual response properties of sSC to V1 we clustered units by following methods laid out in Parker et al., 2023. Briefly, we clustered sSC units with similar gaze-shift responses by performing *k*-means cluster (*k* = 5), resulting in 4 clusters of responsive units and one unresponsive cluster. We used the same clustering metrics when comparing sSC light/dark experiments.

### Detection of burst events

Bursts were defined as periods of spike trains in which a neuron emitted at least three spikes, with an inter-spike interval not exceeding 20 ms and a minimum burst duration of 20 ms. To ensure the robustness and reliability of the data, only neurons exhibiting at least 30 bursts during each recording session were included in the analysis.

### Burst-triggered averages

A burst-triggered average analysis was performed to investigate the temporal relationship between bursts and head/eye movements. In this analysis, angular velocities of both head and eye movements were computed for 20 temporal bins (500 ms) preceding and 40 temporal bins (1 second) following the onset of each burst. These time windows were selected to capture both the immediate and sustained effects of bursts on movement dynamics.

For each recording session, angular velocities of the head and eyes were z-scored to standardize the data across sessions. The cumulative angular velocities of the head and eyes around the bursts were then summed for each temporal bin, generating movement profiles for both the head and eyes. This process was repeated for each individual burst within the session. Finally, the mean and standard error of the mean (s.e.m.) of burst-associated head and eye movements were computed for each neuron, providing a quantitative measure of the neuronal modulation of head and eye movements in relation to burst onset.

### Statistical analysis

Data were statistically analyzed using a custom program written in Python (v 3.8.19). We used the Chi-square test, Wilcoxon rank-sum test, Wilcoxon sign-rank test, Kruskal-Wallis test, and Kolmogorov-Smirnov test to determine statistical significance when appropriate. The Chi-square test was used for proportional comparisons. The Wilcoxon rank-sum test was applied to compare head movement amplitude or TFs/SFs across different behavioral conditions or between groups. We used Wilcoxon sign-rank tests to assess differences in head modulation indices between light and dark conditions. The significance of differences between groups was determined using the Kruskal-Wallis test, Wilcoxon rank-sum test, or Kolmogorov-Smirnov test where normality could not be assumed. Unless otherwise indicated, the significance level was set at α = 0.05. All error bars represent the standard error of the mean (SEM).

## Supplemental Figures

**Figure S1.**
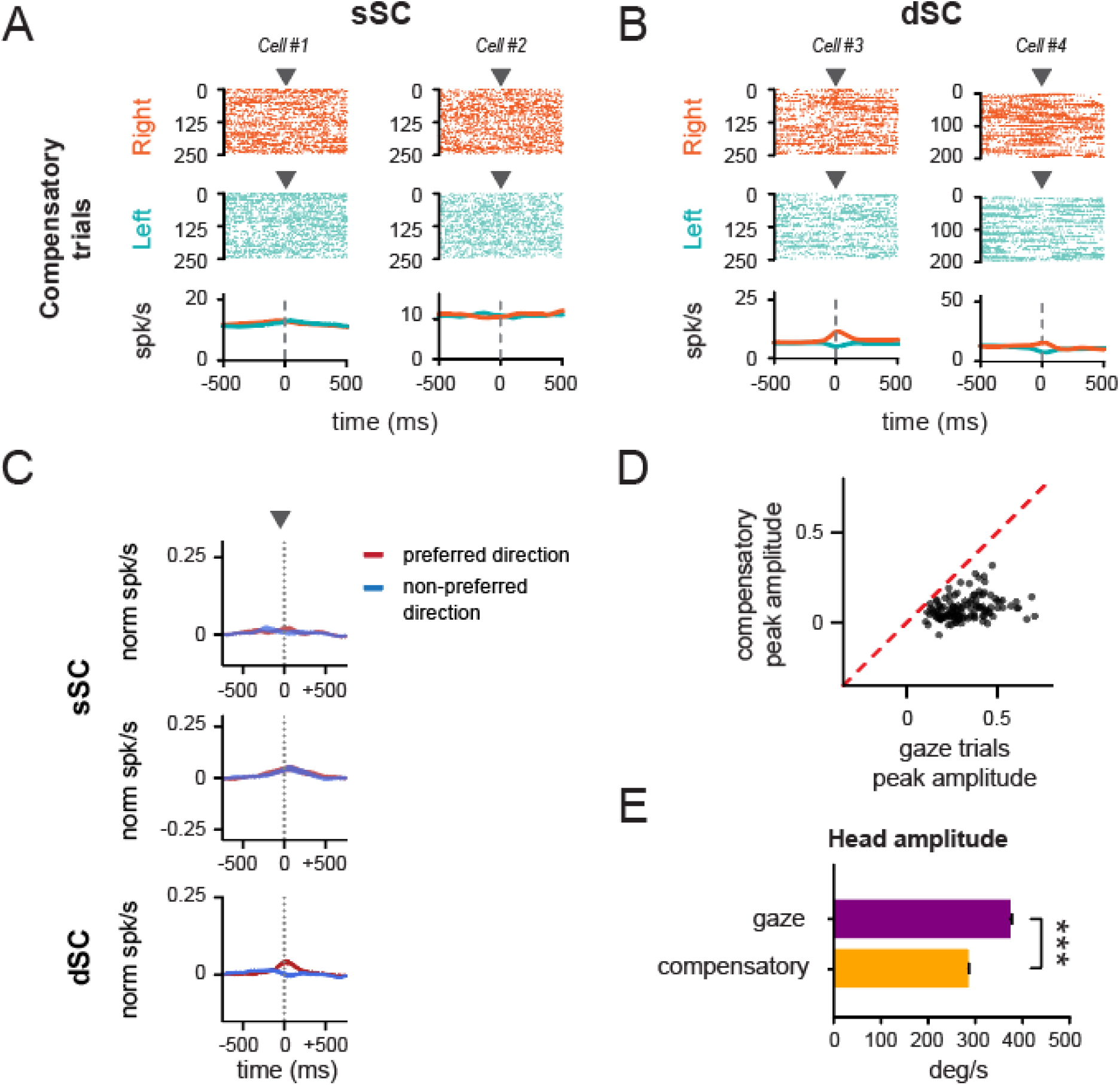
Compensatory responses in SC. (A-B) Trial spike rasters in sSC (*A*) and dSC (*B*) during compensatory movements. Arrowheads indicate onset of the movement. (C) Mean PETHs response for sSC positive responsive cells (*top*), sSC negative responsive cells (*middle*), and dSC responsive cells (*bottom*) during compensatory movements. Red line shows the preferred direction of compensatory response. Blue line shows non-preferred direction of compensatory response. (D) Scatter plot of peak amplitude of responses during gaze and compensatory trials. (E) Bar plot of mean amplitude for gaze and compensatory trials. Error bars indicate s.e.m.

**Figure S2.**
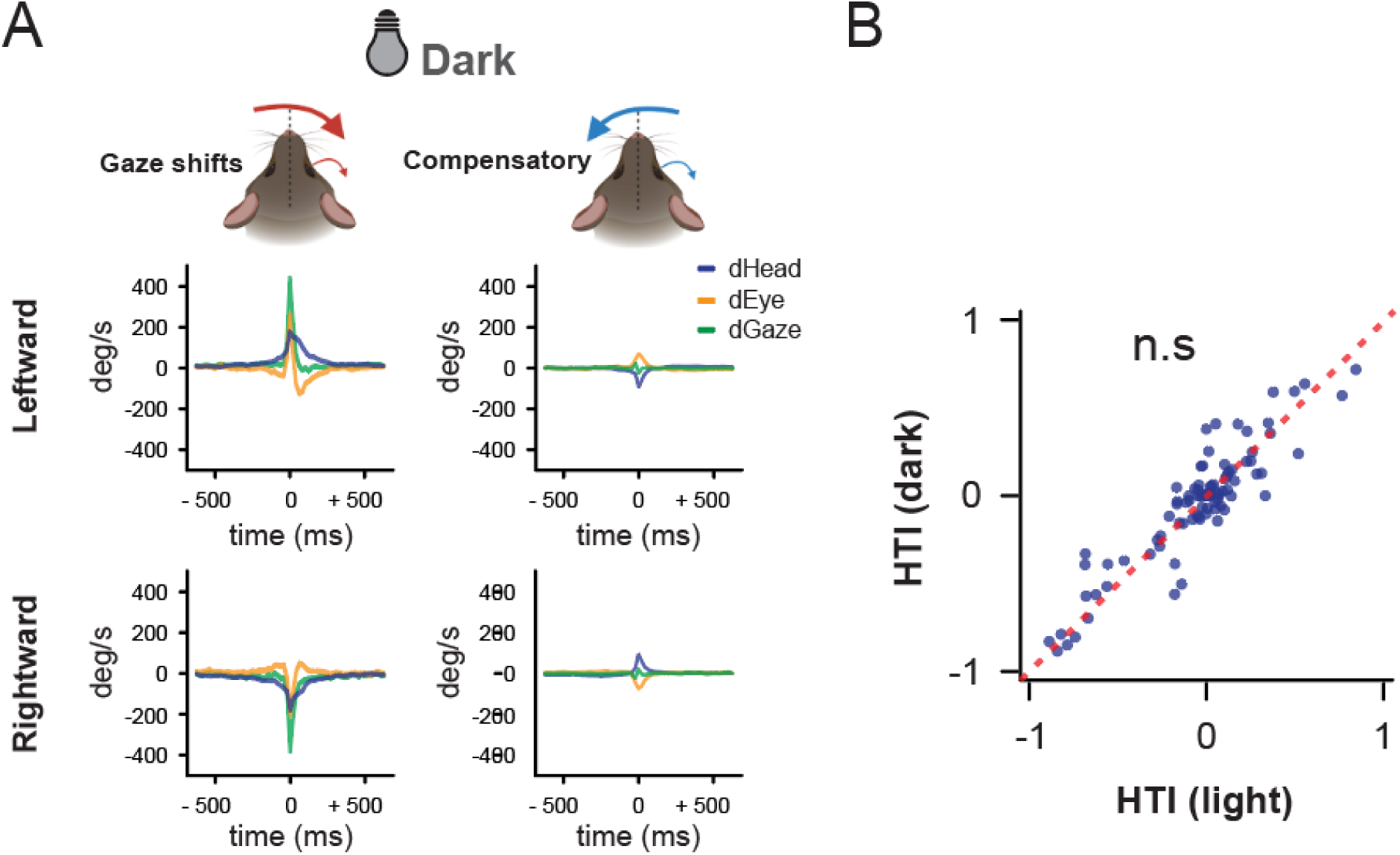
Preserved head tuning in the deep SC. (A) Schematic of head and eye movements in the dark session (n = 6 animals), along with the averaged trace of head, eye, and gaze. (B) HTI distribution from light vs. dark for each cell.

**Figure S3.**
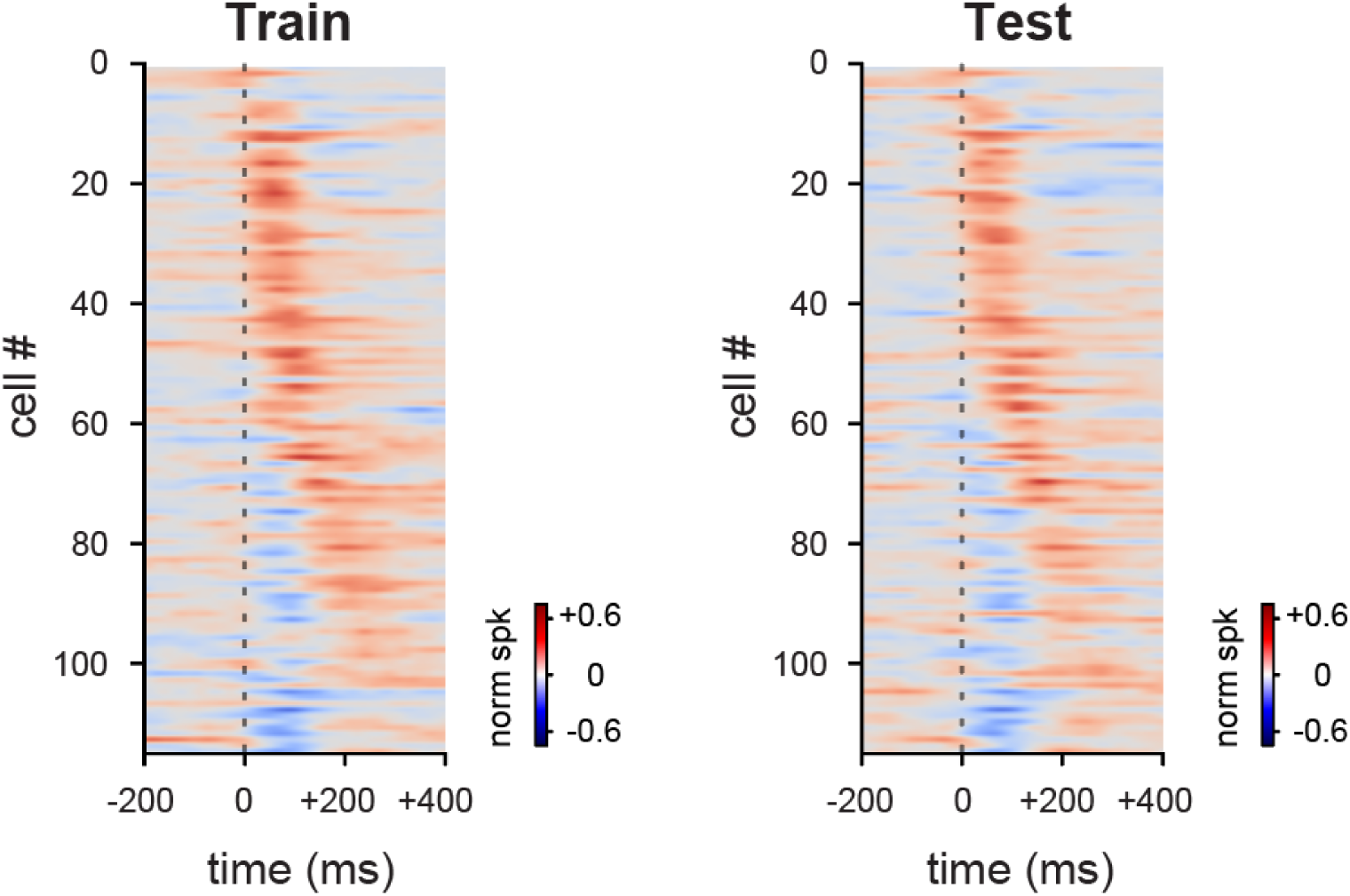
Cross-validation. Cross-validation of gaze shift PETHs of all responsive cells for all responsive cells in sSC. Gaze shift times were randomly divided into two sets used to calculate PETHs in both the train (*left*) and test (*right*) sets. The test set was sorted by the latency of the peak positive response in the train set.

**Figure S4.**
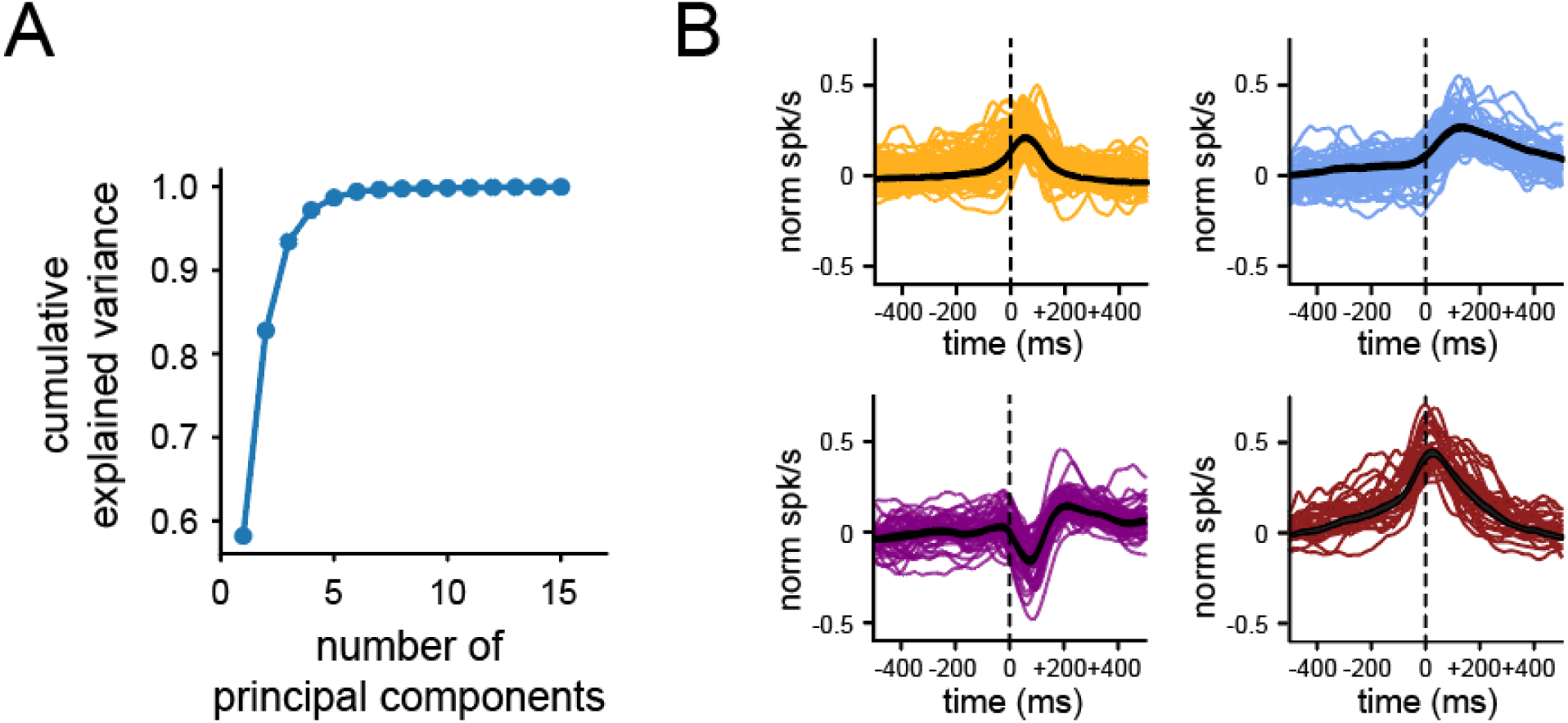
Clustering results when sorted within SC. (A) Elbow plot for PCA showing the cumulative explained variance for each number of principal components. (B) Four types of clustered responses across SC.

**Figure S5.**
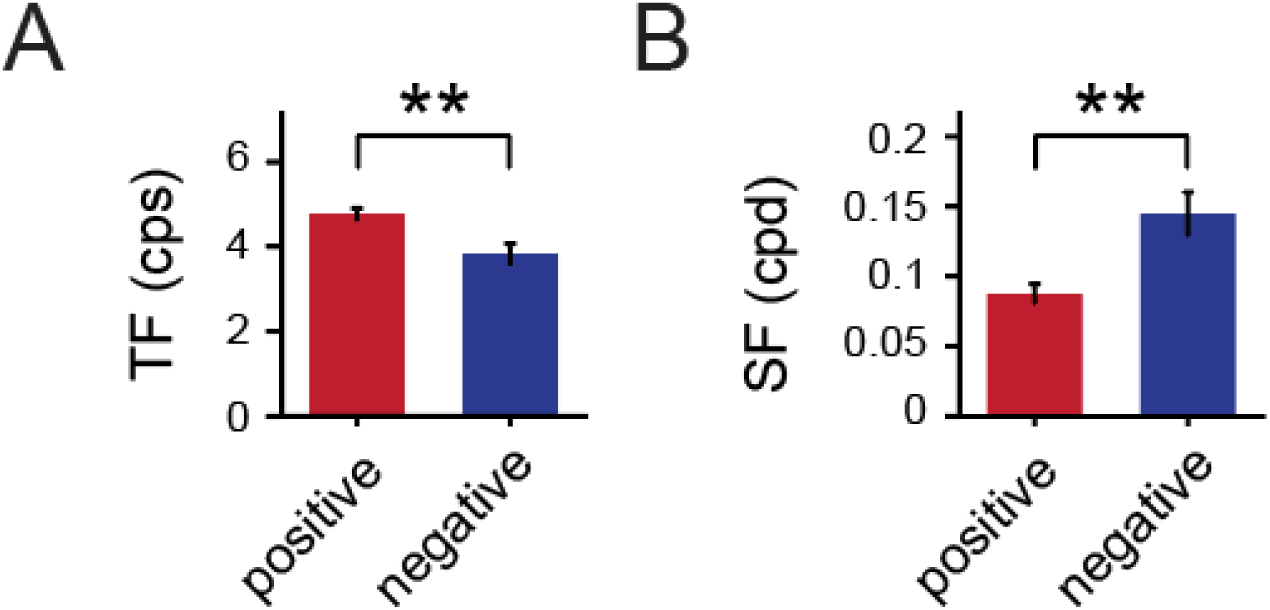
Temporal frequency and spatial frequency preferences across all gaze-responsive positive and negative cells. (A) Mean ± s.e.m. TF tuning preferences for cells in each group (Z = 3.278, *p* = 0.001; Wilcoxon rank-sum test). (B) Mean ± s.e.m. SF tuning preferences for cells in each group (Z = -3.217, *p* = 0.0013; Wilcoxon rank-sum test).

**Figure S6.**
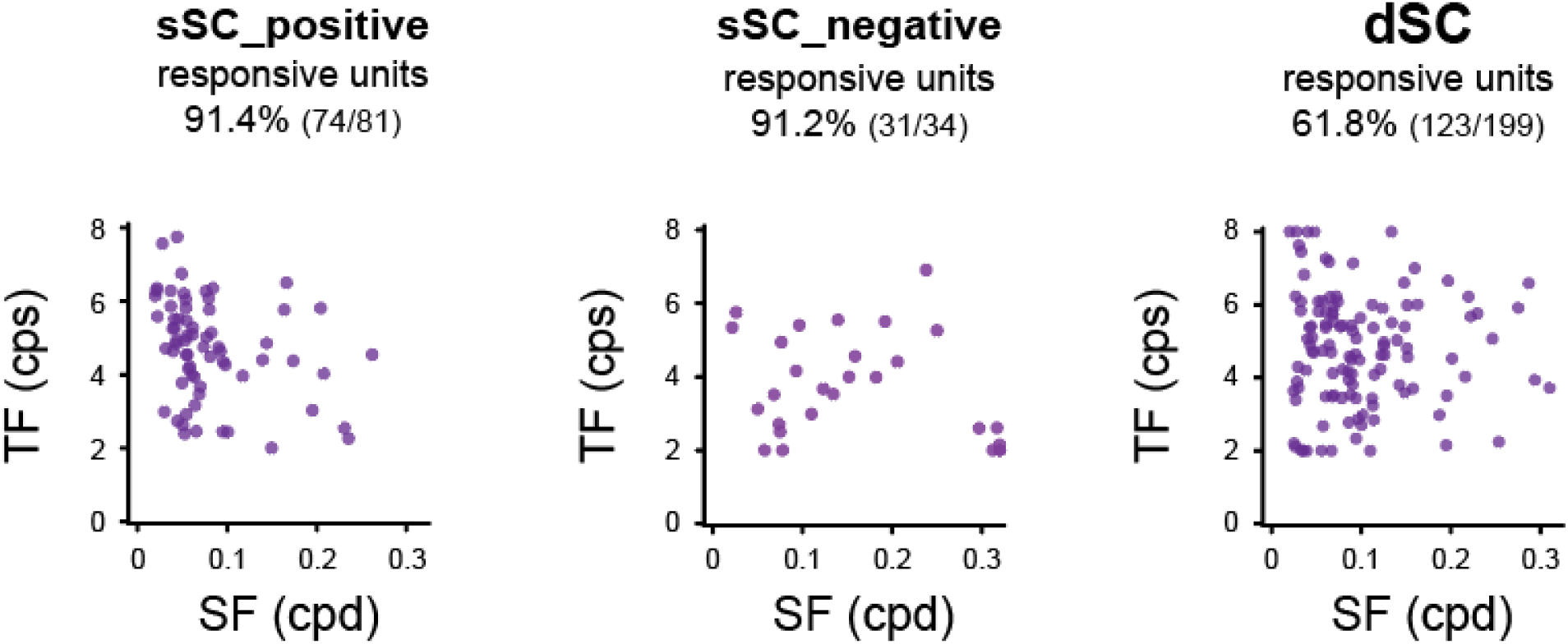
Distribution of SF and TF across the gaze-responsive cells in the SC across different depths.

